# Search calls long duration puts the bat on the tympanate moth's radar. A new perspective on how Noctuoidea's auditory cells drive evasive manoeuvres

**DOI:** 10.1101/037648

**Authors:** Herve Thevenon, Gerit Pfuhl

## Abstract

The auditory stimulation method used in experiments on moth A cell(s) is generally believed to be adequate to characterise the encoding of echolocation signals. The stimulation method hosts, though, several bias. Their compounded effects can explain a range of discrepancies between the reported electrophysiological recordings and significantly alter the current interpretation. To test the hypothesis that the bias may significantly alter our current understanding of the moth’s auditory transducer characteristics, papers using the same auditory stimulation method and reporting on either spiking threshold or spiking activity of the moth’s A cells were analysed. The consistency of the reported data was assessed. A range of corrections issued from best practices and theoretical background were applied to the data in an attempt to re-interpret the data. We found that it is not possible to apply a posteriori corrections to all data and bias. However the corrected data indicates that the A cell’s spiking may be (i) independent of the repetition rate, (ii) maximum when detecting the long and low intensity pulses of the bat in searching mode, and (iii) steadily reduce as the bat closes on the moth. These observations raise the possibility that a fixed action pattern drives the moths’s erratic evasive manoeuvres until the final moment. In depth investigation of the potential bias also suggest that the auditory transducer’s response may be constant for a larger frequency range than thought so far, and provide clues to explain the negative taxis in response to the searching bats’s calls detection.

## Introduction

It has been observed that nocturnal moths apply a range of evasive manoeuvres that appear to depend on the distance of the moth to its main predator: the bat [1]. The likelihood to successfully escape an attack when moths are released 6 m away from a group of foraging bats is 45% [2] irrespective of the moth’s family. This suggests the outcome of the moth’s manoeuvres is not driven by chance. Dissections have shown that moths of the notodontidae family have a single neuron in their tympanal organ - the A cell - [3] that is responsible for the transduction of ultrasounds typically emitted by their foraging predators. Other moth families that are equipped with up to 4 A cells per ear, do not display significant difference in behaviour or escape rate [1].

In anthropomorphic terms, the moth seems to evaluate its distance to the bat by using some characteristics of the bat’s own echolocation pulses. The pulses’ intensity is an intuitive parameter for this evaluation and its effect on the A cell spiking has been studied for more than 50 years. It is well known that the intensity of sound waves decreases with the square of the distance. Therefore the instant evaluation of the intensity of the sounds it receives does not allow the moth to decide whether its predator is far or close. Figure 1 illustrates that from a moth’s perspective a signal received with a given intensity may have been produced by many combinations of intensity of the source and distance to the source.

**Figure 1.**
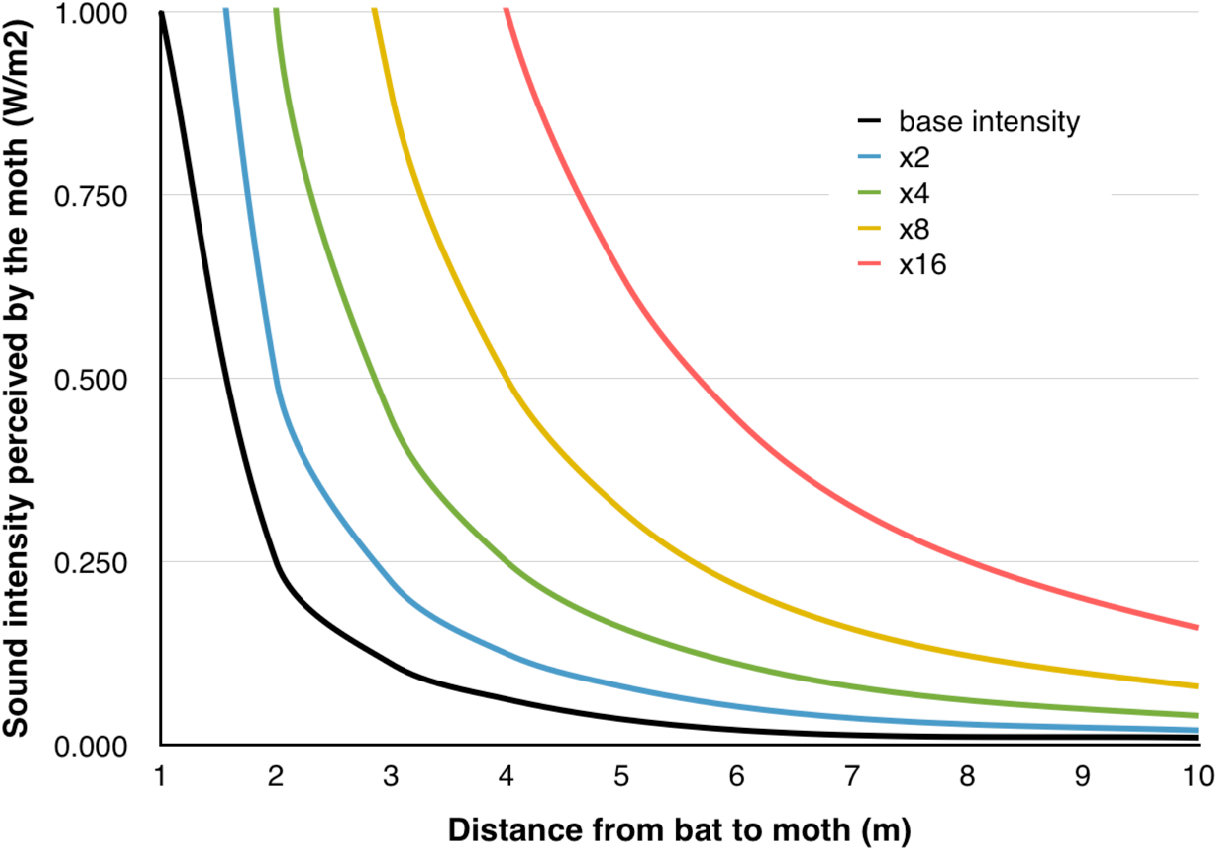
The intensity of the signal received by the moth may have been produced by an infinity of (intensity at source, distance) combinations.

In addition, the ultrasonic pulses of the foraging bat vary during the chase: usually the pulses’ intensity and duration decrease with the distance to the prey whilst pulses are emitted more frequently [4]. Indeed, this observation makes the moth’s intensity-triggered evasive behaviour even more unlikely. It also opens a host of possibilities for the potential parameters involved in the moth’s calculations besides pulse intensity: pulse duration, pulse repetition rate, pulse carrier frequency, and integration over time. The actual combination remains to be determined in order to understand how the complex survival behaviours of the moth are elicited with - for some of them - only one neuron bridging the environment and the nervous system [5].

Since 1957, many experiments have been conducted to characterise the spiking threshold and activity of the nocturnal moths’ A cells. Since the mid 1980s, the same investigation method has been applied to different families and species of moths. This investigation method consists of a moth tethered for extracellular recordings of one of its tympanal nerves whose related ear is stimulated by a sound source emitting ultrasonic pulses. The intensity of the sound received by the moth is controlled by the experimenter by reading the measurement made by a Sound Pressure Level (SPL) meter connected to a microphone (Figure 2).

**Figure 2.**
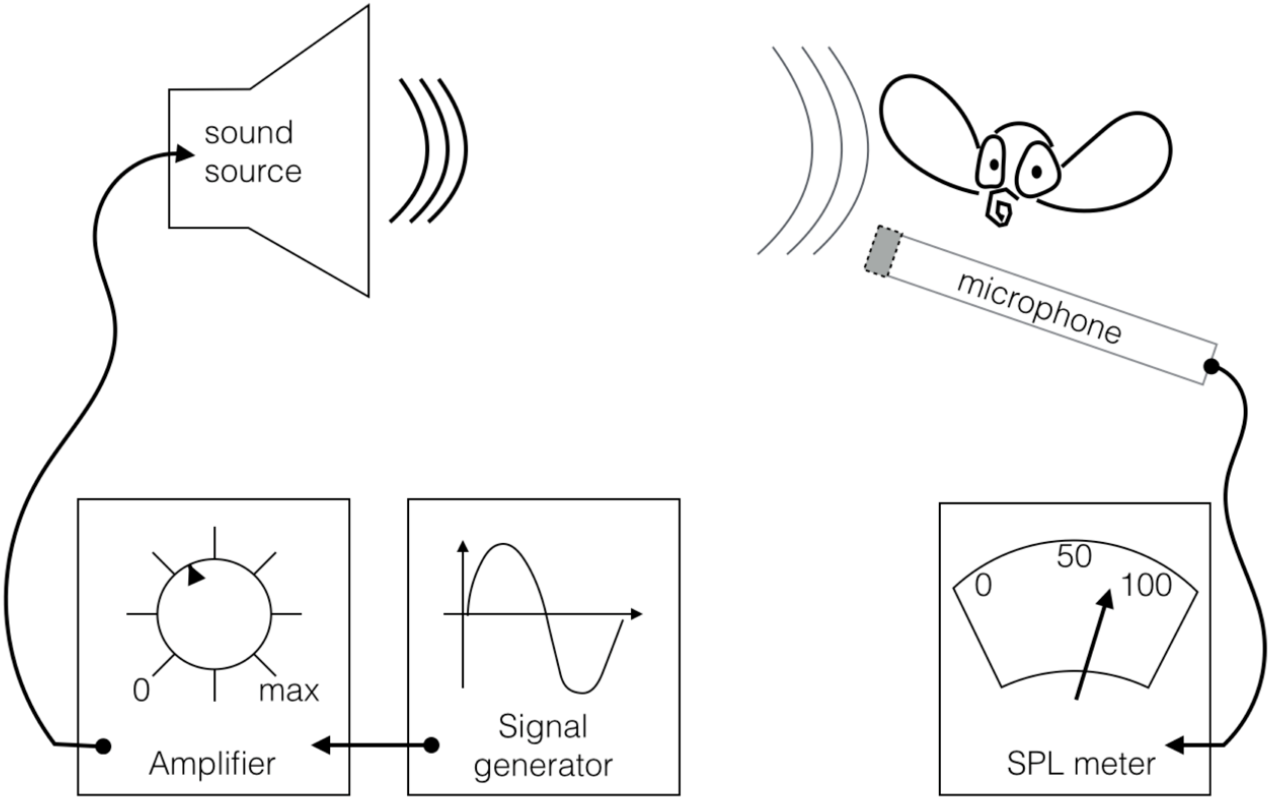
Auditory stimulation setup. A speaker emits ultrasounds towards the tethered moth’s ear. The intensity of the stimulation is controlled via a Sound Pressure Level meter connected to a microphone.

This method has intrinsic biases that we will introduce as required through the course of this paper. The main concern is with the measurement of the sound intensity. What is actually measured via the equipment used in the method is sound pressure, not sound intensity. The rigorous definition of sound intensity is the "vector quantity equal to the product of the sound pressure and the associated fluid particle velocity vector" [6]. It is measured in Watt/m^2^ and describes the flow of energies (e.g. kinetic, potential, heat) through a surface. Sound pressure is a component of sound intensity that is measured in Pascal. Sound pressure may be used as a proxy for sound intensity under specific conditions. As will be seen later, these conditions were not met during the experiments we analysed. This is important because the sound pressure reported is universally used throughout and across experiments using the same method, in order to compare and draw conclusions regarding the characteristics of the moth’s ear as an energy detector [7, 8].

In the absence of technology available to measure sound intensity above 20 kHz [9], microphones remain the most practical tools to evaluate sound intensity in the frequency range that matters to many moths species: 20 kHz ˜ 100 kHz.

The significant variations in the data reported by a series of experiments selected for their methodological similarities were the reasons for us to look into how the auditory stimulation method was implemented. For example, Fullard’s data [10] shows that the SPL required to elicit a given spiking activity increases as the pulse repetition rate increases for pulses of equal duration. Conversely, Waters’s data [11] shows that the SPL required to elicit a given spiking activity decreases as the pulse duration increases at constant repetition rate. The data from both studies are reported in Figures 3 and 4. The energy received by the moth’s ear (over a given period of time) is directly proportional to the pulse duration and repetition rate. Therefore both conclusions and the energy detector hypothesis can not be true simultaneously.

**Figure 3.**
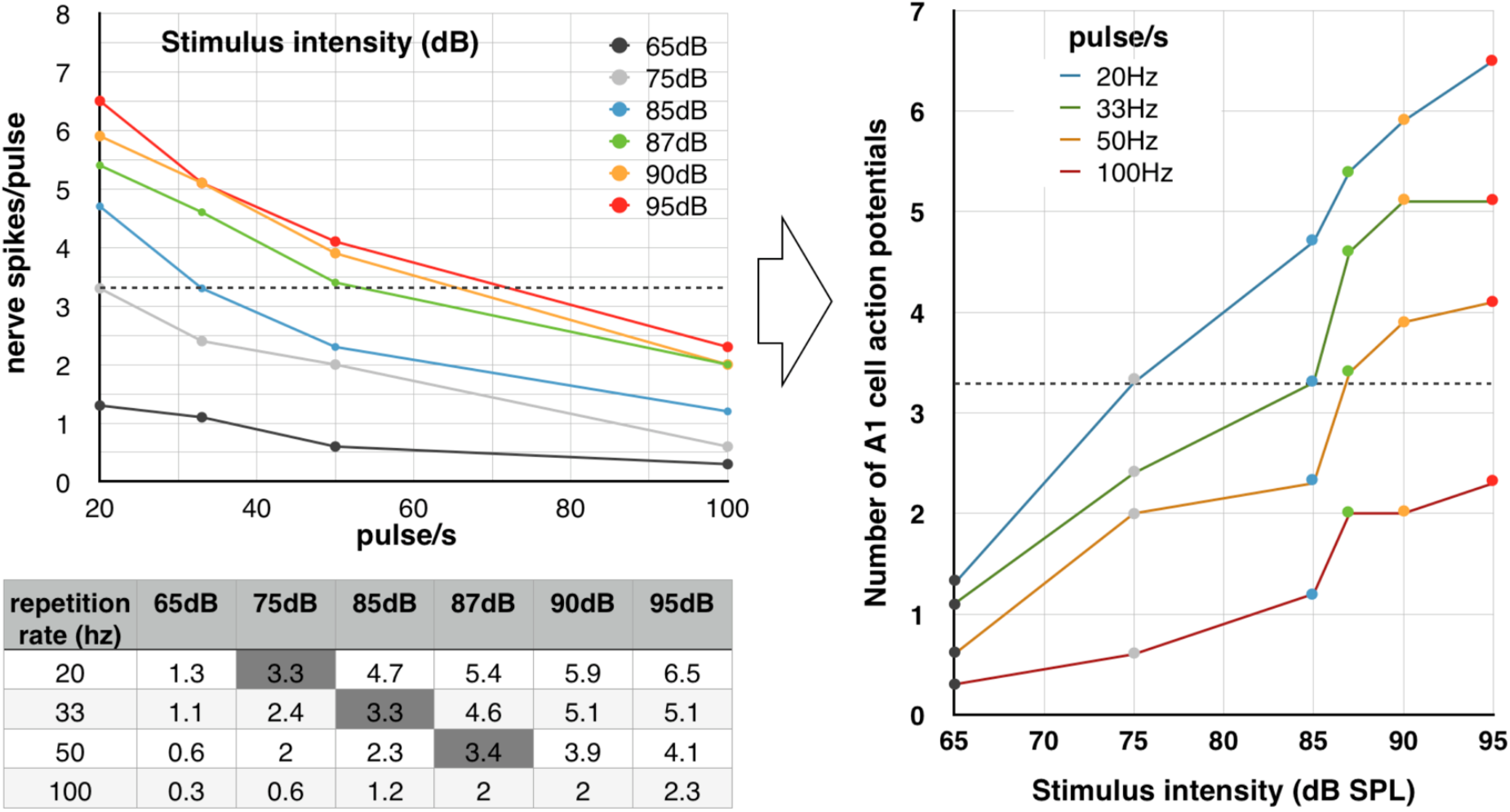
According to Fullard [10], the higher the repetition rate, the more intensity is required to elicit a given spiking activity per stimulus. Top left: graph modified from Fullard. Bottom left: data tabulated. Right: Graph transposed for easier comparison with Figure 4. The horizontal dotted lines indicate an equivalent level of spiking arbitrarily set to 3.3 spikes per pulse.

**Figure 4.**
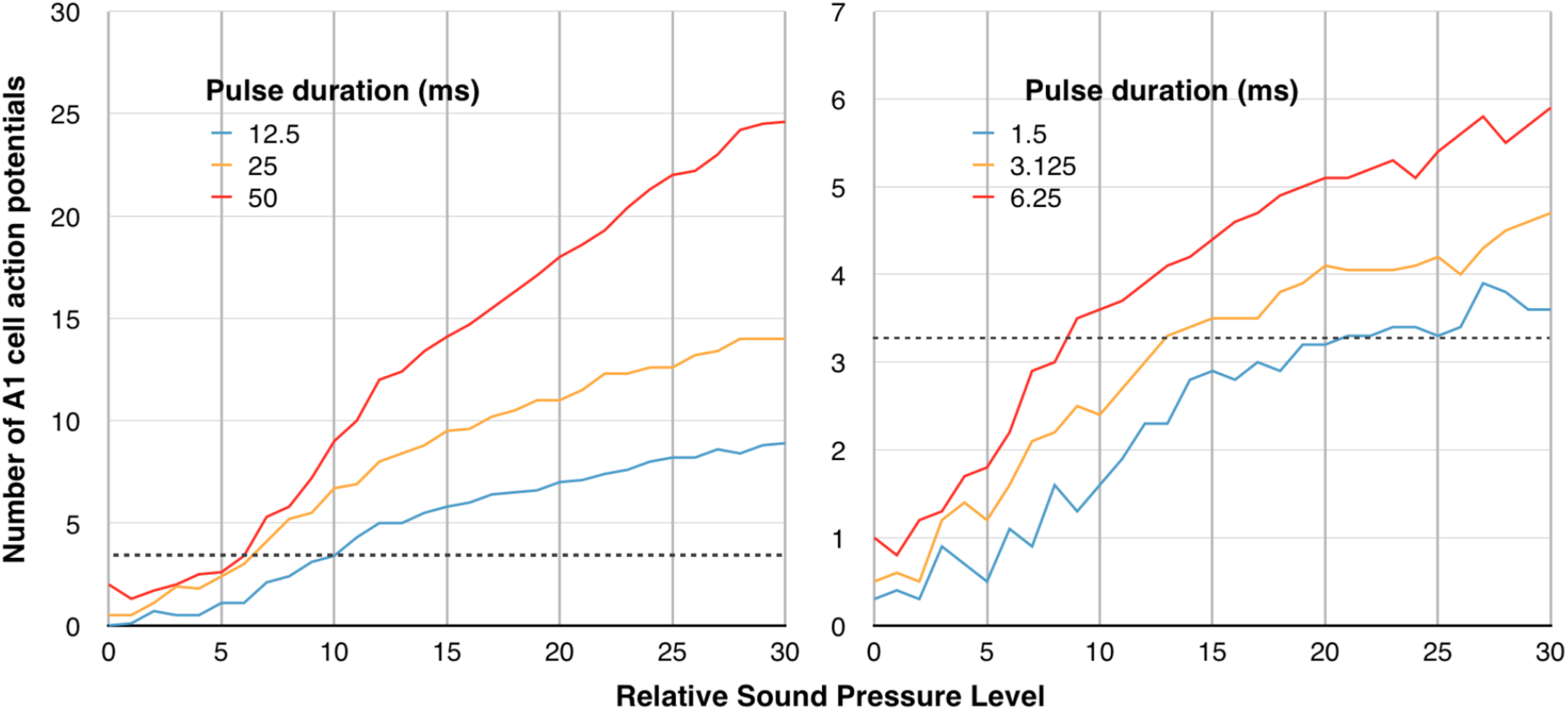
Reproduced from Waters [11]. The longer the pulse duration, the less intensity is required to elicit a given spiking activity per stimulus. Repetition rate is 1Hz. As in Figure 3, the dotted lines indicate an equivalent level of spiking arbitrarily set to 3.3 spikes per pulse.

**Figure 5.**
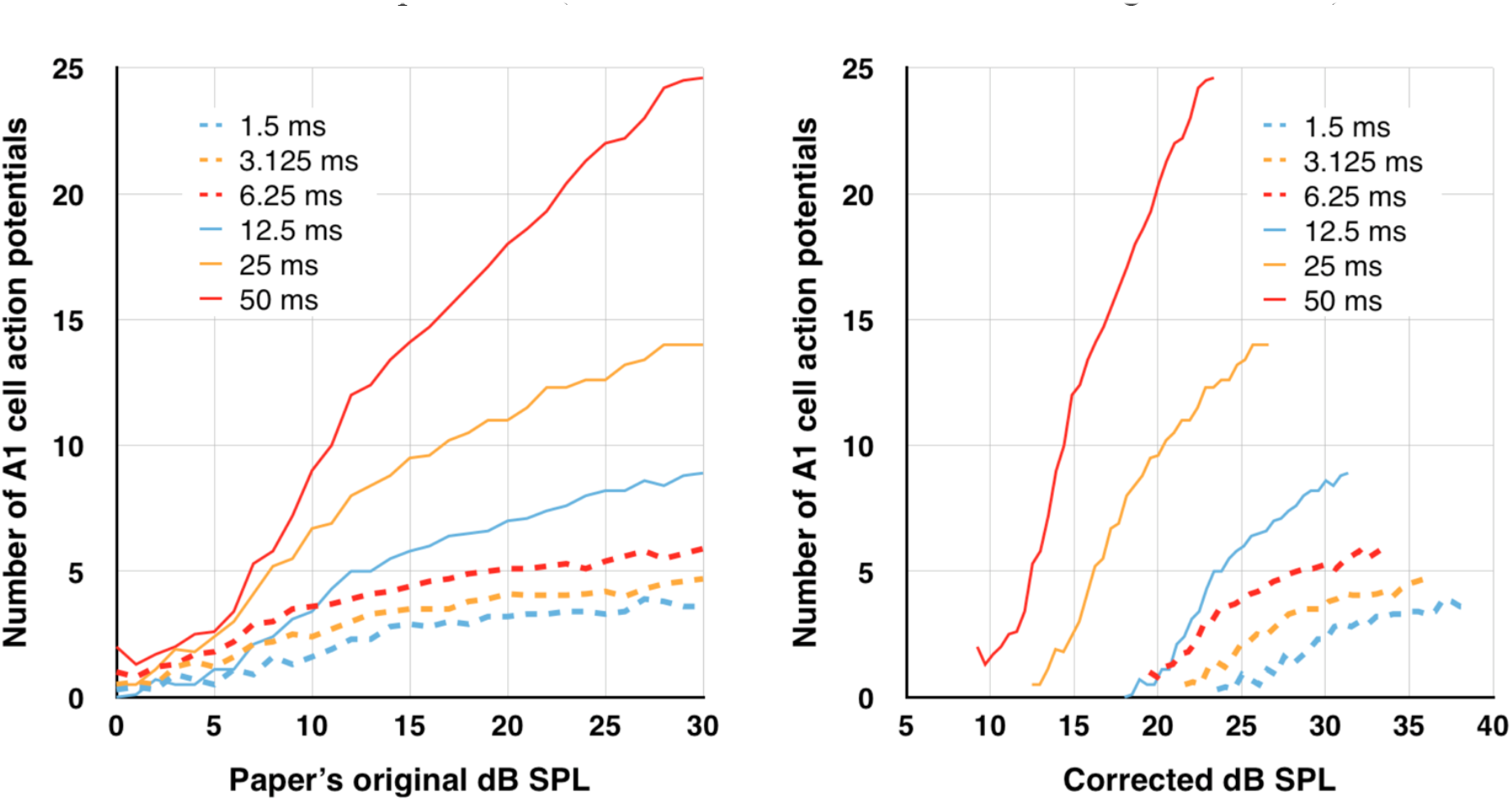
Effect of the correction on Waters’s data. Left: Waters’s original results on the same graph. Right: The same data without the systematic error induced by the damping circuit of the SPL meter.

**Figure 6.**
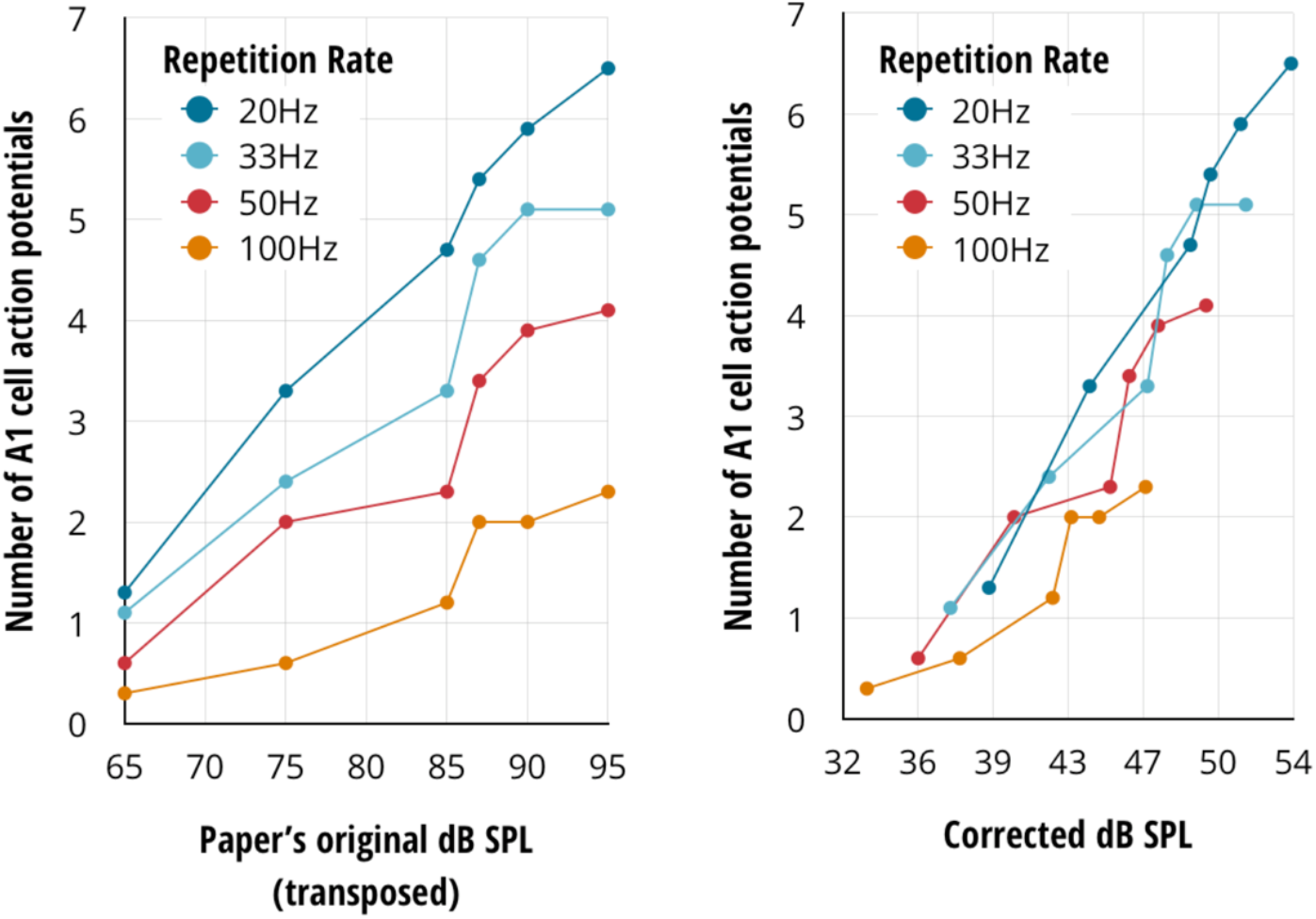
Effect of the correction on Fullard’s data. Left: Fullard’s original data. Right: The same data without the systematic error induced by the damping circuit of the SPL meter.

Here, we provide three results that emerged after scrutinising known biases in the stimulation method, e.g. shortcomings of the Sound Pressure Level (SPL) meter, and applying standard acoustic "corrections". The three results outlined in this paper unfold in cascade. The first result relies heavily on acoustics and theory of operation of the 1960’s SPL meters. This background knowledge allows us to devise a method to infer the actual incident SPL. The second result is derived by building upon the first method by borrowing best practices from sound level measurements as developed in aeronautics. The third result highlights more of the biases found when analysing the properties of the sound stimulation method across all the papers reviewed. It also allows us to formulate an alternate approach.

## Results

**Result 1: Experimental data obtained at fixed frequency may be corrected by inferring the actual SPL received by the microphones. A new conclusion arises from data published by Waters: the A cell firing starts when the bat is in search mode and far from the moth and the firing is maximum. An additional conclusion comes out from one of Fullard’s papers: the firing is independent of the echolocation repetition rate. Taken together, these results suggest the pulse duration is the key to A cell firing.**

For two data sets [10, 11] both using fixed frequency stimulation, we could gauge the actual SPL received by the microphones. These two papers report data about *Agrotis Segetum* - from the Noctuidae family - and *Cycnia tenera* - a member of the Arctiidae family - both belonging to the Noctuiodea super-family. Both species have two A-cells located on the metathorax. In the first instance, we re-analysed data reported by Waters [11]. We refer to the experiment pertaining to the spiking activity of the A1 cell given different pulse duration and the SPL as measured by the SPL meter. The sound stimulation consisted of trapezoidal signals emitted with a constant SPL with a rise/fall duration equal to 10% of a series of plateau durations. The original result stems from two series of experiments (the 3 shortest durations and the 3 longest durations).

Waters’s data corrected for SPL meter’s pulse duration related bias suggests that the A1 cell’s firing rate would be higher when the bat is far away (longer pulses and lower SPL), and would become lower as the bat gets closer (shorter pulses and higher SPL). Waters originally split his experiment in two groups — the shortest durations in one group, and the longest in the other — and reported SPL as relative to the spiking threshold obtained for the shortest duration in each group. Therefore, these relative SPL differ by an unknown offset that is the likely cause for the two groups’ original data to overlap when superimposed. The corrected datasets suggest that more SPL was required to obtain the baseline spiking threshold in the lowest durations group than in the highest durations group, and therefore the lowest durations group’s data should have slightly higher SPL than shown (i.e. should be shifted to the right). With or without this caveat, this result departs from the general view developed through the papers selected. Because inputs with same SPL and different pulse durations produce a different spiking activity, we suggest that the pulse duration is as significant to the auditory transducer’s output as the received SPL.

Fullard [10] investigated the effect of different repetition rates in *Cycnia Tenera.* Since the design of all SPL meters used in this field of research have the same inherent systematic error, we corrected Fullards’ data with the same method used with Waters’s results.

Fullard concluded that the repetition rate was "a cue for defensive behaviour". The corrected data tells the opposite: the spiking frequency is not correlated to the pulse repetition rate when the bias of the SPL meter is taken into account. A linear regression can be applied to the corrected data and reasonably fit the sample of points independently of their respective repetition rate (Figure 7). The residuals did not comply with the Gaussian distribution to back this conclusion, but this may be explained by the small number of samples and their uneven distribution over the SPL new interval.

**Figure 7.**
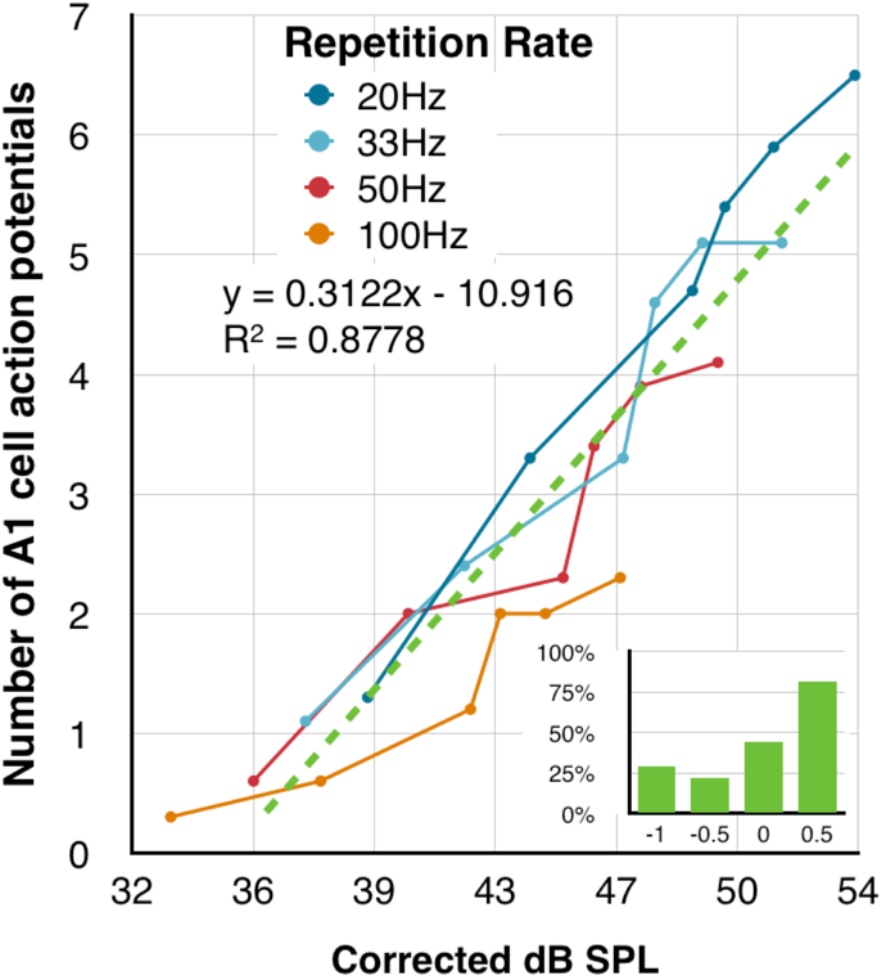
Linear regression applied to Fullard’s corrected data and evaluation of the residuals.

**Result 2: Experimental data obtained while varying frequency are biased by the microphone orientation. A posteriori corrections can not be applied. However, Madsen and Miller’s data and most detailed method suggest that *Barathra Brassicae* (Noctuidae) are insensitive to variations of frequencies in the range 20 kHz ˜ 60 kHz [12].**

The characterisation of the spiking threshold of the A cells was performed across a range of carrier frequencies varying from 20 kHz to 100 kHz. Madsen and Miller reported data on *Barathra Brassicae* (Noctuidae) while using a B&K4138 microphone with grid on. The moth was placed at a 150 degrees angle to the sound source. We can therefore assume that the microphone would have been oriented similarly. Accordingly, we first applied a SPL meter correction and followed with a series of incidence corrections.

The corrected results are presented in Figure 8.

**Figure 8.**
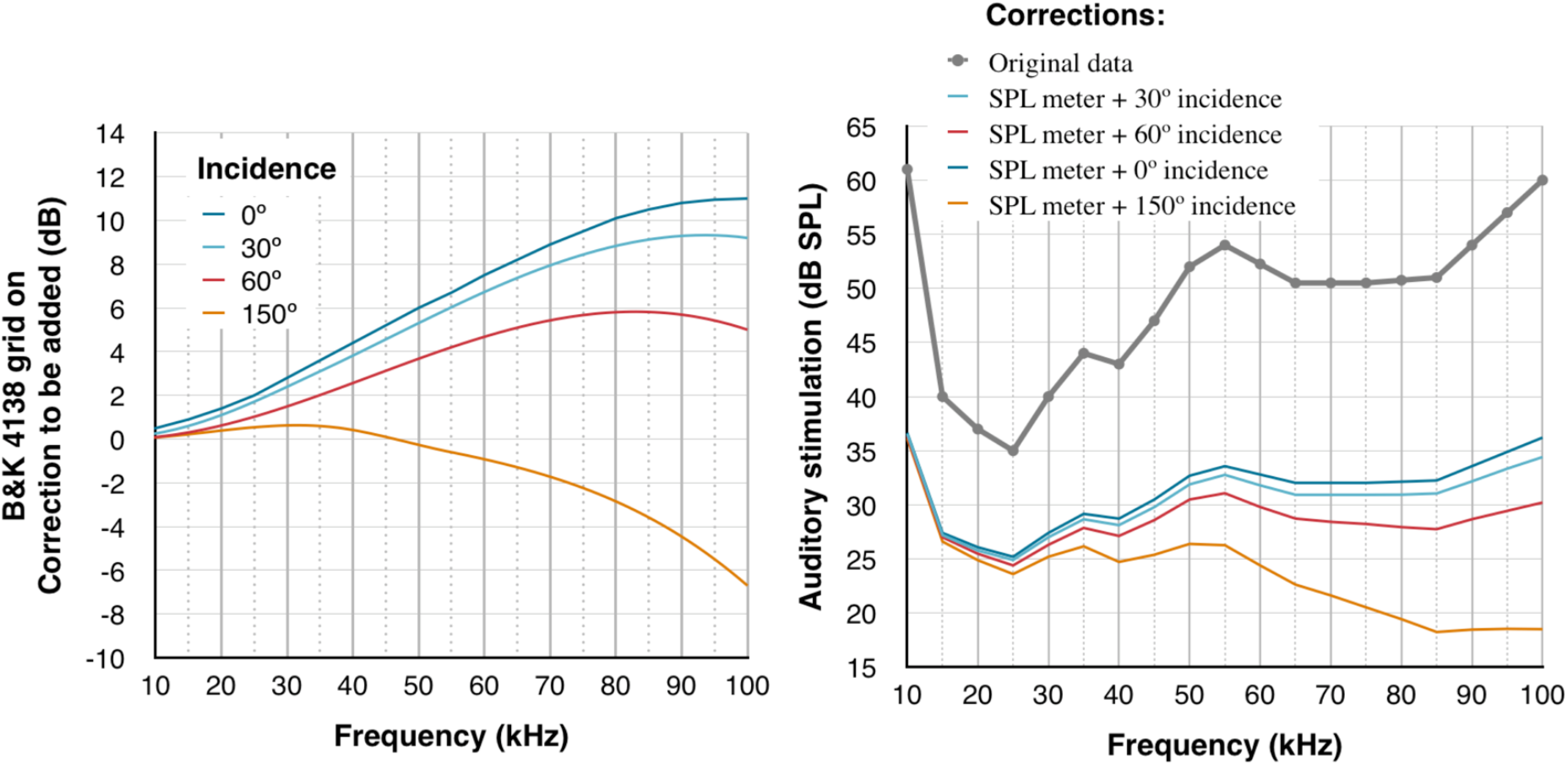
Influence of the microphone orientation on the SPL measurement. Left: Correction of incidence using B & K corrections chart for the 4138 microphone with grid on. Right: The graph shows the original data, the SPL meter bias correction alone, and several microphone incidence corrections compounded with the SPL meter bias correction.

This example shows the importance of the microphone incidence bias when compounded with the SPL meter bias. The correction is frequency dependent and can reach 40dB! The removal of the grid requires applying similar corrections and does not reduce the biases’ impact. None of the papers investigated allowed the application of incidence corrections retrospectively, either because the data are missing, or because the interpretation is ambiguous (see tables 1a and 1b).

**Table 1a.**
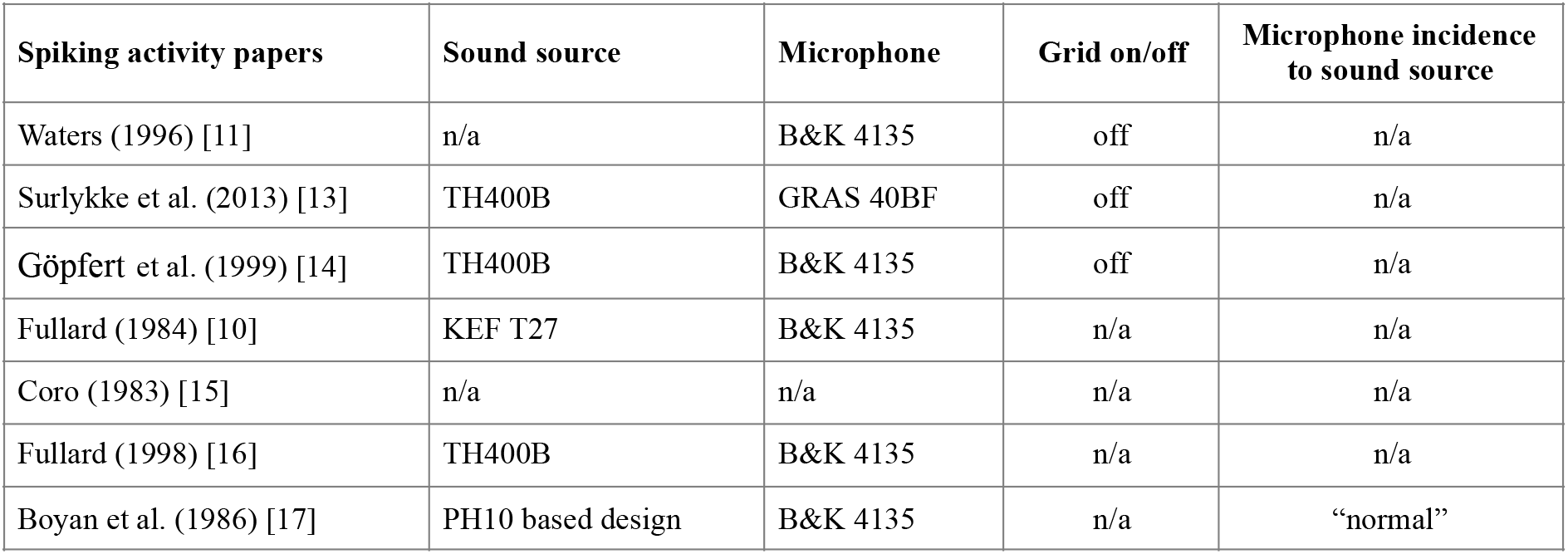
Data related to incidence corrections in the papers focused on spiking activity

| Spiking activity papers | Sound source | Microphone | Grid on/off | Microphone incidence to sound source |
| --- | --- | --- | --- | --- |
| Waters (1996) [11] | n/a | B&K 4135 | off | n/a |
| Surlykke et al. (2013) [13] | TH400B | GRAS 40BF | off | n/a |
| Göpfert et al. (1999) [14] | TH400B | B&K 4135 | off | n/a |
| Fullard (1984) [10] | KEF T27 | B&K 4135 | n/a | n/a |
| Coro (1983) [15] | n/a | n/a | n/a | n/a |
| Fullard (1998) [16] | TH400B | B&K 4135 | n/a | n/a |
| Boyan et al. (1986) [17] | PH10 based design | B&K 4135 | n/a | “normal” |

**Table 1b.** Data related to incidence corrections in the papers focused on spiking threshold

| Spiking threshold papers | Sound source | Microphone | Grid on/off | Microphone incidence to sound source |
| --- | --- | --- | --- | --- |
| Waters et al. (1996) [18] | N/A | B&K 4135 | off | n/a |
| Hofstede et al. (2011) [19] | ScanSpeak 60102 | B&K 4138 | n/a | n/a |
| Madsen et al. (1987) [12] | Bespoke design | B&K 4138 | n/a | “150 degrees” |
| Surlykke et al. (2013) [13] | TH400B | GRAS 40BF | off | n/a |
| Surlykke et al. (2013) [13] | TH400B | B&K 4135 | n/a | n/a |
| Skals et al. (2000) [20] | Bespoke design | B&K 4135 | off | n/a |
| Rydell et al. (1997) [21] | TH400B | B&K 4135 | on | n/a |
| Jackson et al. (2010) [22] | ScanSpeak (no model) | B&K 4135 | n/a | n/a |
| Göpfert et al. (1999) [14] | TH400B | B&K 4135 | off | n/a |

**Result 3: Without meeting free-field conditions and using proper sound sources, all the results are likely to be biased by interference patterns and distorted signal, whose impacts can not be asserted. This collection of biases inherent to the auditory stimulation method leads us to suggest an alternate sound stimulation method.**

Free-field conditions are met when the space around the measuring microphone is exempt of reflected sound waves (including standing waves) and interference. This condition may be met if: (1) there is only one sound source that can stimulate the moth, and (2) the distance between the sound source and any obstacle - including the preparation - is larger than the distance travelled by the sound wave during the pulse duration.

The first condition can be met by shielding the experimentation area against external sound sources in the frequency range 20 kHz ˜ 100 kHz. Anechoic chambers have reportedly been used for this purpose in several experimental setups. Failure to do so means that external sound sources may interfere with the auditory stimulation and contribute to generate spiking for a range of frequencies outside of the stimulation protocol.

Given the second condition and the average propagation speed of sounds in air of 340 m/s, reflections will definitely be avoided if 5 ms pulses are given a 1.65 m clearance and if 30 ms pulses are given 10 meters in every direction. The minimum distance should then be increased accordingly for repeated pulses if the repetition rate is likely to generate interferences at the measurement distance. This places practical constraints on the dimensions of the experimental set-up, that anechoic chambers rated for 20 kHz ˜ 100 kHz range can reduce.

Besides these practical considerations, one should also consider theory. In order to make sure that the measurements are significant, acousticians refer to the far-field or Fraunhofer distance: a distance where the diffraction of the sound waves created by the sound source itself become negligible enough to not create interferences at the distance of measurement. The Fraunhofer distance is defined as twice the sound source’s largest dimension squared divided by the wavelength of the sound wave. Table 2 shows that for all the experiments the preparation was placed too close to the sound source to ensure correct measurements. Free field is defined as a subset of the far field where there is no interference. The standard free field test shows that the intensity of the sound decreases by 6 dB as the distance to the sound source doubles.

**Table 2.** Inventory of the sound sources used in the selected experiments. The table reports physical dimensions, frequency rating, number of times used in the experiments reporting on spiking activity and threshold. ^*^Speakers’ dimension from papers, manufacturer’s documentation or derived from picture (TH400B).

| **Sound sources**<br>Manufacturer and model<br>(alternatively paper reference when custom made) | **Technics TH400B** | **KEF T27** | **Avisoft Scanspeak** | **PH10** | **Madsen & Miller 1986** | **Skals & Surlykke 2000** |
| --- | --- | --- | --- | --- | --- | --- |
| Spiking activity | 3 | 1 | 0 | 1 | 0 | 0 |
| Spiking threshold | 4 | 0 | 2 | 0 | 1 | 1 |
| Rated frequency spectrum | 0~20 kHz | 0~20 kHz | 0~100kHz | N/A | N/A | N/A |
| Largest dimension (mm)* | 70 | 19 | 32 | 5 | 15 | 60 |
| Distance to preparation (mm) | 300 or 400 | 100 | 300 | 4 | 40 | N/A |
| Fraunhofer distance (mm)<br>@100kHz | 2,882 | 212 | 602 | 15 | 132 | 2,118 |

## Discussion

In this paper, we have shown that the conditions required to perform accurate acoustic measurements of the sound pressure levels required to characterise the auditory transducer of the nocturnal moths are never met, and therefore do not allow transposing SPL (dB) to proper sound intensity (W/m^2^). As a result, none of the experiments reviewed could possibly answer the question "Is the moth’s ear an energy detector?" We have seen that sound pressure levels were controlled via a method flawed by several important biases, and that the reported results could contradict one another for the same species. As a countermeasure, we developed a corrective method that enabled reconciling discrepant results for experiments based on a single carrier frequency that may be different between experiments. We also found that we can not apply a posteriori corrections on experiments that perform sound pressure measurements over a range of frequencies, because of three main factors: the actual incidence of the microphone to the sound source is never described in non-ambiguous terms, the distance between the sound source and the preparation is insufficient to prevent interferences in the area of the microphone, and some sound sources are basically inadequate for the task (the TH400B’s diffusion plate effectively creates two distinct sound sources).

The results obtained after applying the corrective method developed in this paper suggest a reevaluation on three levels: (1) A new auditory stimulation method is required; (2) Based on Fullard’s experiments and the novel corrective method presented in this paper, the A cell’s spiking activity seems to be independent of the pulse repetition rate; and, (3) Corrections of Waters’s results suggest that the pulse duration is the key trigger to evoke spiking.

If the new results produced using Fullard’s and Waters’s published data stand the test of experiment, new iterations of the experiments conducted by Madsen and Miller [12] should be conducted to determine whether the moths enter any fixed action patterns after reception of a bat searching call and sustained stimulation with approaching calls. Madsen and Miller noted that "when the flight oscillator is running, auditory stimuli can modulate neuronal responses in different ways depending on some unknown state of the nervous system". From another perspective, the body of experimental data suggests that moths detect searching bats before bats can detect them. As a consequence, experiments like those of Acharya and Fenton’s [2] pertaining to the attack success rate of preying bats on moths released in the vicinity of their foraging area should be extended to larger release distances. We suggest those distances be from 10 to 25 meters away from the foraging area. The rationale is that when released within a 6 meters radius of search calls, without being shielded from those searching calls, the moth’s nervous system may have already modified its internal state. As a result, the current results may not reflect the benefit for the moth to detect a crowd of foraging bats before being detected and subsequently avoid confrontation.

Fullard’s interpretation that the A cell encodes the pulse repetition rate did explain that some moth species exploit the pulse repetition rate to distinguish between bat calls and mating calls [23]. Our interpretation suggests that the exploitation of the pulse rate may take place at the interneuron level [5, 24].

### Basis for a new auditory stimulation method

Auditory stimulations require pre-calibrated sound sources designed to operate in the range 10 kHz ˜ 100 kHz as a whole or for a specific frequency. The sound sources should present a low level of linear and non-linear distortions and be accompanied by manufacturer-supplied free-field correction charts. Pre-calibration must be achieved for the whole sound source system, including the air interface and its pulse/intensity driver.

The sound source and the preparation should be shielded from any other sound source, including any electronic equipment used for the purpose of the experiment. Therefore, the sound source driver should be connected to the sound source by a cable long enough to allow for the driver to be outside the shielding enclosure. Thanks to the preliminary calibration of the sound source and its pulse/ intensity driver, there will be no requirement to insert a microphone within the sound field when the preparation is in the free field space. The use of a microphone in the shielding enclosure should be limited to test for free-field conditions and confirm the sound source calibration and integrity before and after the experiment.

The sound sources may be mono-frequency air transducers. Their small size and diffraction angle weigh positively on the space required to achieve free-field conditions. The sound source and driver should be designed to travel easily between labs and prevent tampering. Ideally, the sound source system can be pre-loaded with a test plan (typically a text file on a removable media) that would allow stimulating the moth automatically and recording the electrophysiology offline. Given the stimulation frequency, the encoded electrophysiological data can be paired with the stimuli by a third party, thus allowing for double-blind experiments.

Since there is virtually no sound intensity probe that operates above 10 kHz, the experiments require free-field conditions. This may be achieved with a large cylindrical chamber that will enclose the sound source and preparation only, shielding them from any external sound source. Roeder and Treat [25] should be noted for presenting a similar set-up, that seemed to borrow from waveguide theory. We suggest making the cylinder long and wide enough to minimise internal reflections. This can be planned by using the sound source’s manufacturer specifications. Length will mainly depend on the size of the sound source. It should be several multiples of the most conservative Fraunhofer distance in order to allow for testing free field conditions within the cylinder. The sound source should protrude slightly from one extremity of the cylinder. The other extremity of the cylinder should be prepared with anechoic structure and materials suitable to absorb sound waves in the range 10 kHz ˜ 100 kHz. If such material cannot be found or is too expensive to source, the shield will need to be extended in order to prevent reflections that might interfere with the incoming pulses from the source in the area of the preparation.

The preparation should be installed in the midline of the cylinder using fittings whose size will not diffract the wave plane and create unwanted interferences. A thin gauge (sub-wavelength diameter) taut cable may be used for this purpose. Electrode wires should run longitudinally as to avoid reflections that could interfere around the preparation. Whether the set-up is adequate can be determined using the canonical free-field test: 6dB decreases are observed as the distance doubles.

## Materials and Methods

### Literature selection

First, we searched for papers containing the keywords "moth" or "lepidoptera" and "auditory" or "acoustic" in Scopus, ISI Web of Science then checked all the references for additional materials. In order to be selected, the papers had to report original data for one or both of the following experiments conducted on the nocturnal moths: (1) spiking frequency as a function of carrier frequency, pulse duration, and sound pressure level; and, (2) spiking threshold as a function of sound pressure level, pulse frequency and pulse duration. This is a natural choice when one considers that the spiking frequency and threshold are the most basic characteristics that may be studied in this context, and therefore are the most studied. Such results can only be compared if the experimental conditions and the auditory stimulation method are well described. Consequently, the selected papers provide specific details about the auditory stimulation method applied: pulse duration, pulse shape, pulse repetition rate, frequencies being used. As all the papers presented the data as graphs, the selection process discarded those papers whose data could not be tabulated, typically because units or scale were missing. In order to assess the bias specific to experiments involving ultrasounds, the papers retained include most of the following details: the make and model of the sound source, microphone and SPL meter, distance and incidence of the sound source to the preparation, incidence of the microphone with regards to the sound source. The final selection consists of 12 papers published between 1983 and 2011 that encompass a range of 10 moth species and six moth families. Seven papers [11, 13-17] reported on spiking activity, specifically the number of spikes obtained with stimuli varying in sound pressure level and pulse duration at a fixed frequency. Nine papers [12-14, 18-22] reported on spiking threshold, specifically the pulse frequency and sound pressure level required for a stimuli of a given pulse duration to elicit spiking threshold. However, as will become clear below, for correcting for the systematic bias of the SPL meter, we could only use two studies that used similar experimental conditions on closely related species [10, 11]. In order to take into account the incidence error in microphones, only one study provided enough details [12]. The remaining papers had insufficient details to carry out the corrections (see tables 1a and 1b). Nevertheless, given similar peripheral processing across insect taxa (Hildebrandt et al. 2015) differences between tympanate moth species should be of minor importance given our focus.

### Systematic bias of SPL meters

Late 1960’s SPL meters were not suitable for controlling the intensity of a pulsed sound source. Essentially, these SPL meters were galvanometers enhanced with a time-weighting damping circuit-applied to the signal received from the microphone used as input - to facilitate the readings. The effect of the analogue electronic circuit used for this damping can be expressed in mathematical terms [26] as equation 1.

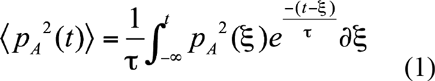

p_A_ is the sound pressure (in Pascal), τ is the damping time constant (1 s for the S setting and 0.125 s for the F setting), ξ is the integration variable, and t is the time of the observation. The older the signal, the less significant.

SPL meter reports dB SPL. dB SPL are obtained by dividing the square of the measured sound pressure by the square of sound pressure of reference (20 micro Pascal). In order to compute equation (1), we expressed it as a Riemann sum shown as the numerator in equation (2). This expression factors out the constants found in equation 1, and by convention ksi is replaced by x. Equation (2) was coded to calculate and graph the movements of the SPL meter’s pointer over time as the meter receives pulses.

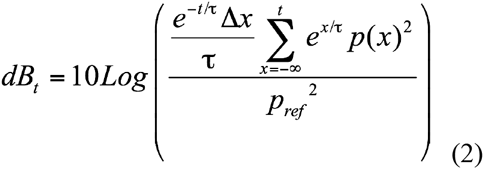

In practice, the -∞ boundary in equation (2) must be replaced by a more computable value "z". Drawing from Marsh’s considerations [26] we found that z = −5 ms is computationally efficient and representative of _—∞_ for any of the trapezoidal signals used in the reminder of the paper.

Figure 9 presents an example of these calculations for pulses whose duration is either 5 ms or 50 ms, and whose repetition rate is either 1 Hz or 10 Hz. For all eight scenarios, the intensity of the signals is arbitrarily set to 60 (j,Pa - i.e. 3 times the SPL of reference, or 9.5 dB SPL. The figure clearly illustrates the difference between the S and F settings. On the F setting, the graph indicates that the pointer jets across the display with a speed that makes the readings impractical. Since the SPL meter setting was never documented in our selection of papers, we assume that the S setting was used, and we used this setting for all the results presented in this paper.

**Figure 9.**
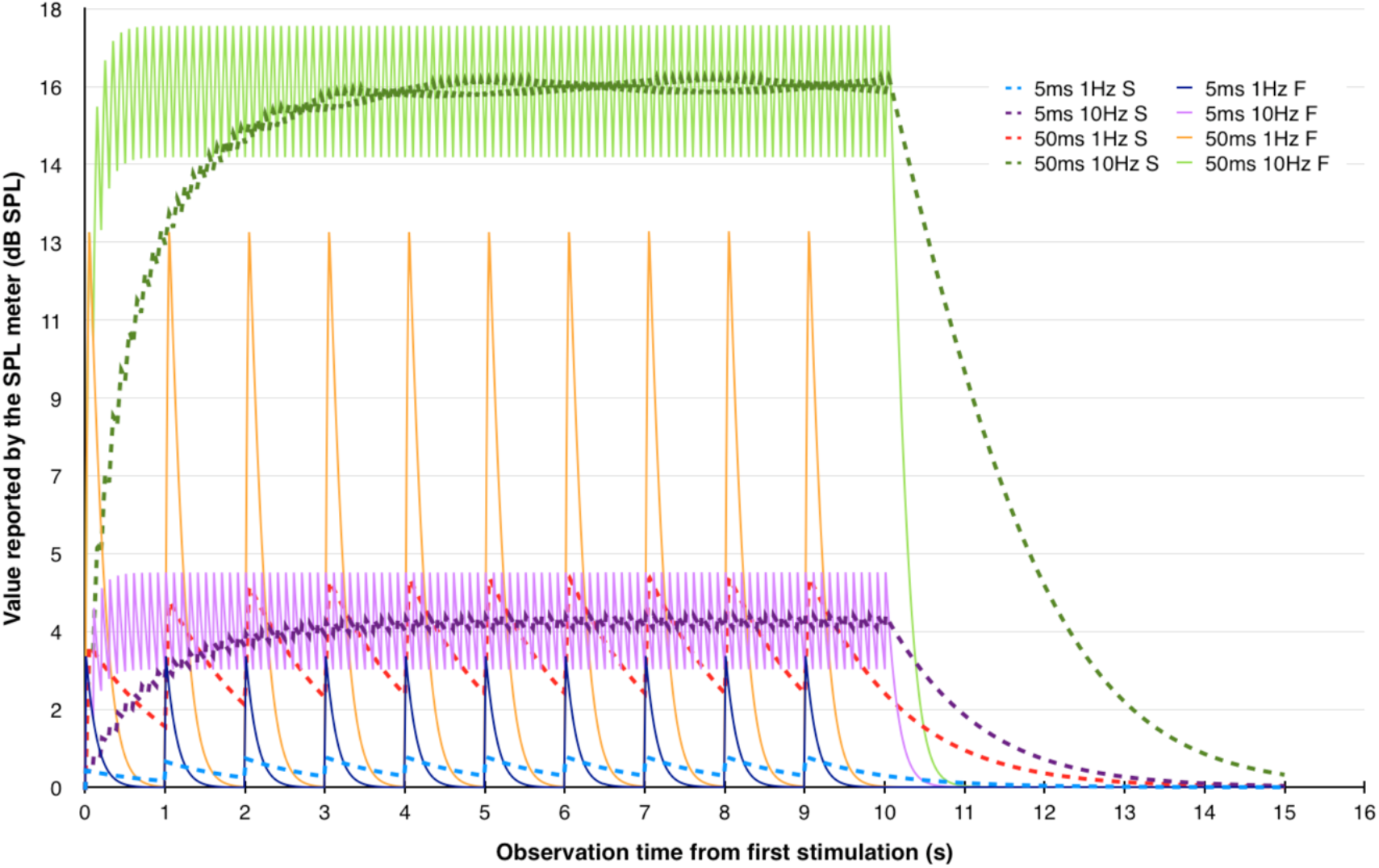
***Graph of the SPL meter’s pointer movements*** for 5ms and 50ms signals, repeated once or 10 times per second, evaluated using the S and F settings.

On this basis, it is possible to infer the actual sound pressure level that would have been returned by a SPL meter complying with equation (1) for a series of measurements that have in common 3 of the 4 following parameters: (i) pulse shape and duration, (ii) pulse carrier frequency, (iii) pulse repetition rate, (iv) measured dB SPL. The method consists in 3 steps:

1. Simulate the response of the SPL meter for the kind of pulses used in the experiment and for a series of emitted sound pressure levels,
2. Find a suitable regression that allows converting the RMS dB SPL readings into the sound pressure level that was received by the SPL meter’s microphone,
3. Replace the RMS dB SPL value as reported in the papers by the corresponding dB SPL value obtained via the regression formula,

We sampled the response of the mathematical SPL meter for signals of different sound pressures for each different plateau duration for Water’s (1996) experiment. For each plateau duration we extracted a regression rule that allowed us to infer the emitted sound pressure level that matches the experimenter’s reading. Figure 10 shows two examples for plateau durations equal to 50 ms and 6.25 ms respectively. Figures 5 and 6 are the outcome of steps 3.

**Figure 10.**
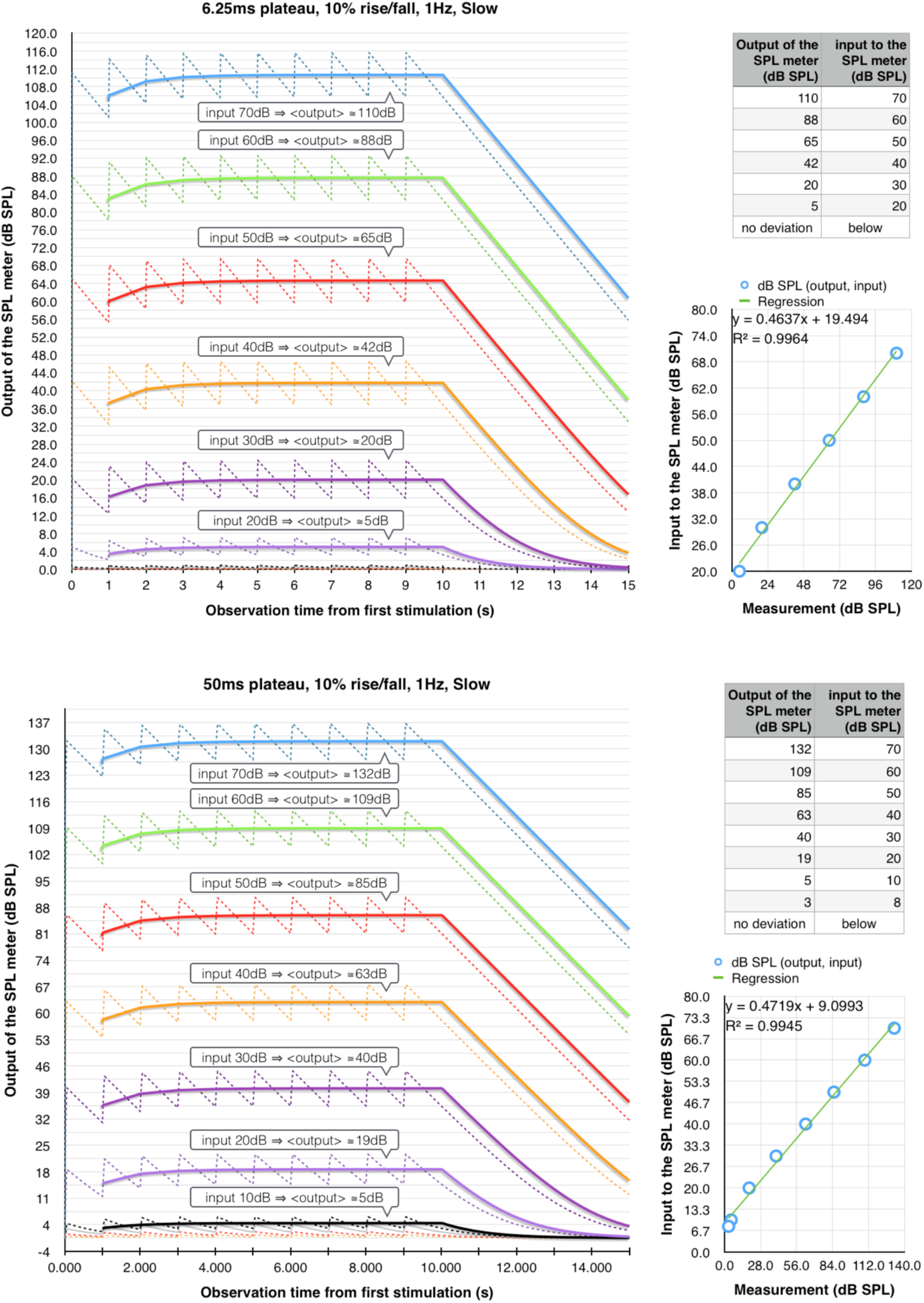
Illustration of steps 1 and 2. Step 1: For each plateau duration, sample the emitted sound pressure space and derive the RMS dB SPL value calculated by the SPL meter according to equation 1. Step 2: Find a suitable regression that converts RMS readings to the actual emitted sound pressure level.

The bias impacts all the papers reviewed (table 3a and 3b).

**Table 3a.** Papers reviewed regarding spiking activity.

| Spiking activity papers | Pulse duration (ms) | Repetition rate (Hz) | SPL meter | SPL meter design |
| --- | --- | --- | --- | --- |
| Waters 1996 | 1.5, 3, 6.25, 12.5, 25, 50 | 1 | B&K2204 | 1969 |
| Surlykke 2003 | 10 | 5 | ? | N/A |
| Gopfert Wasserthal 1999 | 30 | 10 | B&K2606 | 1969 |
| Fullard 1984 | 3 | 1 | ? | N/A |
| Coro 1983 | 45 | 1 | ? | N/A |
| Fullard 1998 | 45 | ? | B&K2607 | 1970 |
| Boyan 1986 | 10 | 1 | B&K2606 | 1969 |

**Table 3b.** Papers reviewed regarding spiking threshold.

| Spiking threshold papers | Pulse duration (ms) | Repetition rate (Hz) | SPL meter | SPL meter design |
| --- | --- | --- | --- | --- |
| Waters & Jones 1996 | 10 | 1 | B&K2204 | 1969 |
| Hofstede and al. 2011 | 20 | 5 | ? | N/A |
| Madsen Miller 1986 | 10 | 10 | B&K2606 | 1969 |
| Surlykke and al 2003 | 10 | 1 | ? | N/A |
| Surlykke and al 2003 | 5 to 30 | 1 | ? | N/A |
| Skals and Surlykke 2000 | 50 | ? | B&K2607 | 1970 |
| Rydell, Skals, Surlykke, Svensson 1997 | 10 | 1 | B&K2606 | 1969 |
| Jackson, Asi, Fullard 2010 | 20 | 1 | B&K2610 | <1980 |
| Gopfert Wasserthal 1999 | 30 | 3 | B&K2331 | ? |

### Microphone incidence error

Microphones are sensitive to the incidence of the sound waves on the membrane and the grid, and this sensitivity varies with the signals’ carrier frequency. The linearity of the microphone’s response is guaranteed for normal incidence and grid on. Therefore, if the microphone’s incidence deviates from 0 degrees, the correction consists of two steps. First, one must find the actuator response for the given frequency by removing the 0 degrees correction for normal incidence and then add up the correction related to the reported degrees incidence [27]. In Madsen and Miller’s paper, the incidence seems to be 150 degrees. Since a SPL meter was involved to report the sound pressure level, the same correction method applied for Fullard’s and Waters’s data should indeed be applied on Madsen’s and Miller’s data. But which correction should be applied first? The diagram in Figure 11 explains the reasoning: the SPL meter’s displayed measure dB2 is based on the microphone’s output dB_1_.

**Figure 11.**
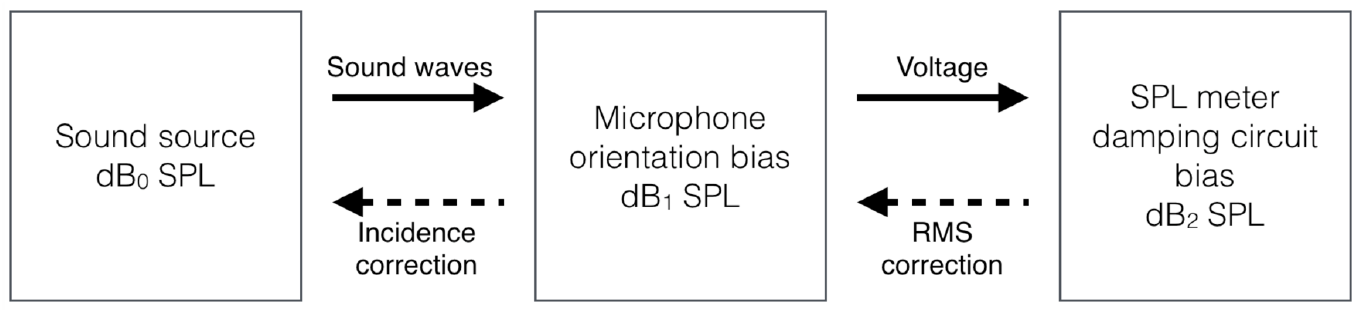
Compounded errors and order of corrections.

Therefore dB_2_ can be expressed as dB_2_ = E_2_(dB_1_) = E_d_(E_i_(dB_0_)) with E_d_ and E_i_ the respective damping and incidence errors induced by the equipment.

Accordingly, we simulated the SPL response to trapezoidal signal of 10 ms including a 0.5 ms raise/ fall time repeated 10 times per second emitted with different sound pressure levels along with a linear regression describing the transformation dB_1_ = T(dB_2_) derived. Subsequently, a series of incidence corrections were applied by using the principle presented above and the manufacturer’s specifications.

## Acknowledgements

We acknowledge the help of Dr Tove Dahl and Dr Ian Welch for their proofreading.

